# Phase-variable restriction–modification systems dynamically regulate plasmid transmission in *Neisseria gonorrhoeae*

**DOI:** 10.64898/2026.01.10.698759

**Authors:** Tabea A. Elsener, Ana Cehovin, Wearn-Xin Yee, Aden Forrow, Sam Palmer, Christoph M. Tang

**Affiliations:** Sir William Dunn School of Pathology, South Parks Road, University of Oxford, Oxford OX1 3RE, United Kingdom; Mathematical Institute, University of Oxford, Andrew Wiles Building, Radcliffe Observatory Quarter, Oxford OX2 6GG, United Kingdom

## Abstract

Horizontal gene transfer (HGT) in bacteria is shaped by restriction–modification systems (RMSs) that define genetic barriers between species and lineages. *Neisseria gonorrhoeae* harbours numerous RMS, most of which are part of its core genome. Here, we show that phase variation in these RMSs can limit interspecies and intraspecies plasmid exchange. While NlaIV likely influenced the acquisition of pConj from the meningococcus, phase variation of the NgoAV RMS produces epigenetically heterogeneous populations that present variable bottlenecks to within-species plasmid transmission, influencing the spread of anti-microbial-resistance plasmids, pConj and p*bla*. A LEAT sequence extends or retracts a molecular ruler within the HsdS specificity subunit, altering spacing of recognition motifs, while truncation of the protein leads to a switch from non-palindromic to palindromic recognition motifs. Thus, while the repertoire of gonococcal RMSs is largely conserved, phase-variation dynamically modulates gene flow, shaping plasmid transmission, and the evolution and spread of antimicrobial resistance.

## INTRODUCTION

Horizontal gene transfer (HGT) facilitates the rapid evolution of bacteria by providing novel genetic traits, with resistance plasmids being major drivers of the spread of antimicrobial resistance (AMR)^1^. However, not all outcomes of HGT are favourable. Plasmid carriage can impose a burden on host bacteria^2^, while the acquisition of heterologous DNA can result in less fit gene variants, and phage infection can be fatal. To mitigate the risks of HGT, bacteria have evolved an array of defence systems that limit HGT^3^, with restriction-modification systems (RMSs) being the most abundant; over 80% of bacteria harbour at least one RMS^4^.

RMSs include a methyltransferase (MTase), which methylates host DNA in a sequence-specific manner, protecting it from the activity of the endonuclease (REase) of RMSs. REases cleave non-methylated foreign DNA^5^, protecting bacteria from phage infection and acquisition of heterologous DNA, including plasmids during transformation or conjugation^6,7^.

RMSs are classified into four types^8^. Type I RMS are multi-enzyme complexes consisting of an HsdM MTase, HsdR REase, and an HsdS specificity subunit. Type I MTases are formed by a HsdM_2_S_1_ complex^9^, and REases are a complex of HsdR_2_M_2_S_1_^10^. The target recognition domains (TRDs) of HsdSs mediate sequence-specific methylation and restriction by type I systems^11,12^. Type II RMSs consist of individual MTases and REases, while the MTase of type III RMSs consists of two MTase subunits; addition of two REase subunits results in the restriction complex^13^. Type IV RMSs consist solely of REases that recognise and cleave unmodified DNA^14^.

The repertoire and activity of RMSs in bacteria generate specific patterns of DNA methylation and regulate gene flow between species and lineages^15,16^. The effect of RMSs can differ depending on the mode of DNA transfer^17,18^, and by the specific combination of RMS and plasmid, including the number and location of restriction sites on a plasmid, the rate of plasmid transfer, and the presence of plasmid-encoded anti-restriction factors^19^. Therefore, understanding the impact of RMSs on the spread of plasmid-mediated resistance requires detailed knowledge of the efficiency of RMSs in preventing plasmid uptake, and plasmid and RMS diversity within a bacterial population.

*Neisseria gonorrhoeae* causes a sexually transmitted infection and is a WHO priority pathogen due to the emergence of AMR^20,21^. Chromosomally-mediated resistance in the gonococcus often arises through incorporation of homologous sequences from other *Neisseria* species^22^. Compared with other Gram-negative priority pathogens, only two AMR plasmids, pConj and p*bla*, are found in the gonococcus, which have contributed to the discontinuation of tetracycline and penicillin therapy^23^. pConj has been acquired by the gonococcus from *Neisseria meningitidis* on more than one occasion^24^, and undermines the use of doxycycline post-exposure prophylaxis (Doxy-PEP) to prevent gonococcal disease^25^. This plasmid is remarkably mobile, with conjugation rates of up to 80% for certain pConj variants in isogenic matings^24,26^. p*bla* is mobilised by pConj but at lower frequencies, with p*bla* often co-resident with pConj^27,28^.

Remarkably, *N. gonorrhoeae* isolates harbour between 13 and 16 RMSs^29,30^, while other non-transformable bacteria with comparable genome sizes usually contain two or three systems^31^. Interestingly, RMSs in the gonococcus can be phase-variable, with their expression predicted to undergo ON:OFF switching or change sequence specificity through alterations in repetitive DNA sequences^30,32^. Furthermore, some phase-variable type I and type III MTases have also been implicated in epigenetic regulation of gonococcal virulence and antibiotic resistance^33,34^.

pConj and p*bla* are unevenly distributed in the population^27,35^, even though pConj is highly mobile, suggesting there are barriers to its spread in the gonococcus. Methylation of *Escherichia coli* plasmids can enhance their transfer into *N. gonorrhoeae*^36^, indicating that RMSs might prevent the acquisition of plasmids by the gonococcus. However, the extent to which RMSs modulate plasmid spread between populations of *N. gonorrhoeae* is unknown.

Here, we examined the distribution and diversity of RMSs across 3,761 *N. gonorrhoeae* whole-genome sequences, with the aim of identifying systems that influence the distribution of resistance plasmids. We found that most RMS loci are part of the gonococcal core genome, distinct from *N. meningitidis*^16^. Through logistic regression analysis of RMSs including their phase-variable features and plasmid distribution, we identified three RMSs associated with plasmid presence/absence. We show that these gonococcal RMSs differ in their efficiency to prevent p*bla* and pConj transfer. The NlaIV endonuclease is highly effective at preventing plasmid acquisition from donors lacking an active NlaIV methylase. As most gonococcal isolates have an active NlaIV MTase, this system is likely to limit the acquisition of plasmids from other species and restrict the plasmid repertoire in this important pathogen. In contrast, we show that variation in the HsdS of the type I NgoAV RMS affects spacing between sequence motifs, with the protein acting as a molecular ruler, which is extended and retracted by a repeat LEAT sequence. HsdS can also switch between a monomeric protein with two distinct TRDs (TRD1 and 2) and a dimeric version which forms an MTase with a unique HsdM_2_S_2_ stoichiometry; this switch mediates a change from modification of non-palindromic to palindromic DNA sequences; loss of NgoAV activity increases plasmid acquisition, consistent with logistic regression. However, plasmid transfer rates between non-isogenic isolates reveal that other factors, beyond RMSs, influence plasmid flow across the gonococcal population.

## EXPERIMENTAL PROCEDURES

### Bacterial strains and growth

Strains used in this study are listed in **Supplementary Table 1**. *N*. *gonorrhoeae* was grown on gonococcal base media (GCB) with 1% v/v Vitox (Oxoid) and 1.1% w/v Bacteriological Agar No. 1 (Oxoid). In liquid, *N*. *gonorrhoeae* was grown in GCB (GCBL) with 1% Vitox and incubated at 37°C in 5% CO_2_ with shaking at 180 r.p.m.. As needed, media were supplemented with kanamycin (kan) at 50 μg/ml, carbenicillin (carb) at 2.5 μg/ml, tetracycline (tet) at 2 μg/ml, erythromycin (ery) at 1 μg/ml (all Sigma), and 6.4% w/v 2-deoxygalactose (2-DOG, Sigma)^37^.

### Transformation of gonococci

*N. gonorrhoeae* was transformed as previously^38^. In brief, 1 μg of DNA in water was spotted on GCB plates and allowed to dry. *N. gonorrhoeae* grown for 16 – 18h on GCB was streaked over the spots and incubated for 8h at 37°C in 5% CO_2_. Bacteria were plated onto selective GCB plates and incubated for 24–72h. Transformants were re-streaked on GCB plates containing antibiotics.

### Strain construction

Primers used in this study are listed in **Supplementary Table 2**. *ngoAV* and *nlaIV* were modified using markerless gene editing^37^. Target genes were replaced with *aph(3)-galK*; the cassette was subsequently replaced by the desired construct, using 2-DOG counter-selection; FA1090 *hsdS* was exchanged with *hsdS* from isolates 60755, NG102, FA1090 and 44569 (**Supplementary Table 1**). To generate isogenic FA1090 strains with *nlaIV_E*_OFF_, FA1090 *nlaIV_E* was amplified from FA1090, and an additional T was introduced at the end of the *nlaIV_E* tract. Strains with *nlaIV_M* modifications were derived from FA1090 *nlaIV_E*_OFF_. To limit phase variation of *nlaIV_M*, the poly-A tract was interrupted by a synonymous A to G substitution at nt. 6 of the tract; removal of an A from the 3’ end of the tract resulted in the *nlaIV_M*_OFF_ strain.

For generating FA1090 NEIS2765 isogenic strains, full-length NEIS2765 was amplified from isolate 60755 (**Supplementary Table 1**), fused with *aph(3)* using Gibson assembly and transformed into FA1090. All strains were confirmed by Sanger sequencing.

To distinguish between donor and recipients in the conjugation assays, strains were marked with *ΔpilD::ermC* and *ΔpilD::aph* constructs, respectively^27^. *pilD::ermC* donors carrying pConj.1 and p*bla*.1 were obtained by conjugation with FA1090 *pilD::aph(3)* pConj.1 p*bla*.1 for 6h, before selection on ery/tet/carb^27^. Individual colonies were then picked, and *porB* sequenced to confirm strains.

### Conjugation assays

Plasmid transfer between isolates was quantified as previously^27^. In summary, donor (*pilD::ermC*) and recipient (*pilD::aph*) strains were grown overnight on GCB agar, sub-cultured in 5 ml GCBL / 1% Vitox to mid-exponential phase, and the bacterial density was adjusted to 10^8^ CFU/ml. Donor and recipient strains were mixed in a 10:1 ratio, and bacteria (5 μl) were spotted onto GCB agar. After 6h, bacteria were harvested into 200 μl GCBL, serially diluted and plated on GCB agar with the appropriate antibiotics (ery for donor only, kan for recipient only, kan/tet for pConj transconjugants and kan/carb for p*bla* transconjugants). Plasmid transfer frequencies were defined as the number of transconjugants/recipient, and experiments performed on four occasions.

### Analysis of whole genome, RMS, and plasmid sequences

Whole genome sequence data from 3,761 *N. gonorrhoeae* isolates (Supplementary Table 3) available on PubMLST (https://pubmlst.org/neisseria)^39^ were analysed. Genetic relationships between the gonococci were determined using *N. gonorrhoeae* core genome multi-locus sequence typing scheme (Ng_cgMLST) version 1.0^40^ and visualised with Grapetree^41^, with a threshold of 400 allelic differences to define core genome clusters (Ng_cgc_400_) ^40^.

RMSs sequences were identified in PubMLST^39^ using the Genome Comparator tool and BLAST with reference sequences^30^. Loci were manually curated, and gene absence was validated by Artemis^42,43^. A nucleotide sequence database of all RMS loci and associated alleles defined in PubMLST was generated, organised into an “RMS in *Neisseria gonorrhoeae*” scheme, and verified by Genome Comparator.

To define repeat lengths, multiple sequence alignments of alleles were generated with Clustal Omega^44^. For loci where simple sequence repeats were detected, features such as the number of nucleotides in a polynucleotide tract, the number of nucleotide repeats, or the position of stop codons were annotated.

### Logistic regression

For each of the 33 RMS loci examined, we extracted features expected to affect RMS behaviour, such as whether a mutation had induced a premature stop codon. We then fit logistic regressions predicting whether pConj or p*bla* were present in each isolate. Each logistic regression had a sample size of 3,761 isolates. Logistic regression for each feature generated a regression coefficient, indicating how strongly the feature contributed to plasmid presence, and p-value for a null model where isolates are independent, as an approximate measure of uncertainty. As isolates are not independent, the coefficients and p-values cannot be used directly to estimate causal effects of features. Instead, we used the product of the p-value and regression coefficient to identify candidate RMSs.

### DNA methylation

DNA from FA1090 with different *hsdS* variants was sequenced using Nanopore (v14 library prep, R10.4.1 flow cells, PlasmidSaurus) to detect adenine methylation. Reads were aligned to the FA1090 genome (PubMLST id: 2855) using Dorado^45^, and sorted and indexed with Samtools^46^. Genome-wide base modifications and predicted motifs were called with Modkit^47^ pileup and motif search.

### Statistical analyses

Data was analysed in R version 4.1.1 using base R and the tidyverse package^48^. Plots were generated with ggplot^49^. Statistical significance was assessed using the Welch two-sample t-test or ANOVA with Tukey multiple comparisons.

## RESULTS

### Association of RMS features and plasmid carriage in the gonococcal population

To examine relationships between the two AMR plasmids, pConj and p*bla*, and RMSs, we initially defined the repertoire and variation of RMSs in the gonococcal population and annotated RMS loci in a dataset of 3,761 isolates obtained from 56 countries between 1936 and 2019 (**Supplementary Table 3**). We identified 16 RMSs in our dataset, encoded by 33 loci (**Supplementary Table 4**), with type II RMSs accounting for 12 RMSs, and type I (n=2) and type III (n=2) RMSs also present. Of note, 28 of the 33 RMS loci are part of the core gonococcal genome, *i.e.* present in ≥95% of isolates in the population. Indicating that the repertoire of RMS is conserved in this species. We therefore constructed the gonococcal population structure based only on their RMS allelic differences and found it was strikingly similar to that obtained with 1,668 core genes (using the clustering system Ng_cgc_400_, **Figure 1A**), highlighting that genetic variation of RMSs mirrors variation in the remainder of the genome.

**Figure 1:**
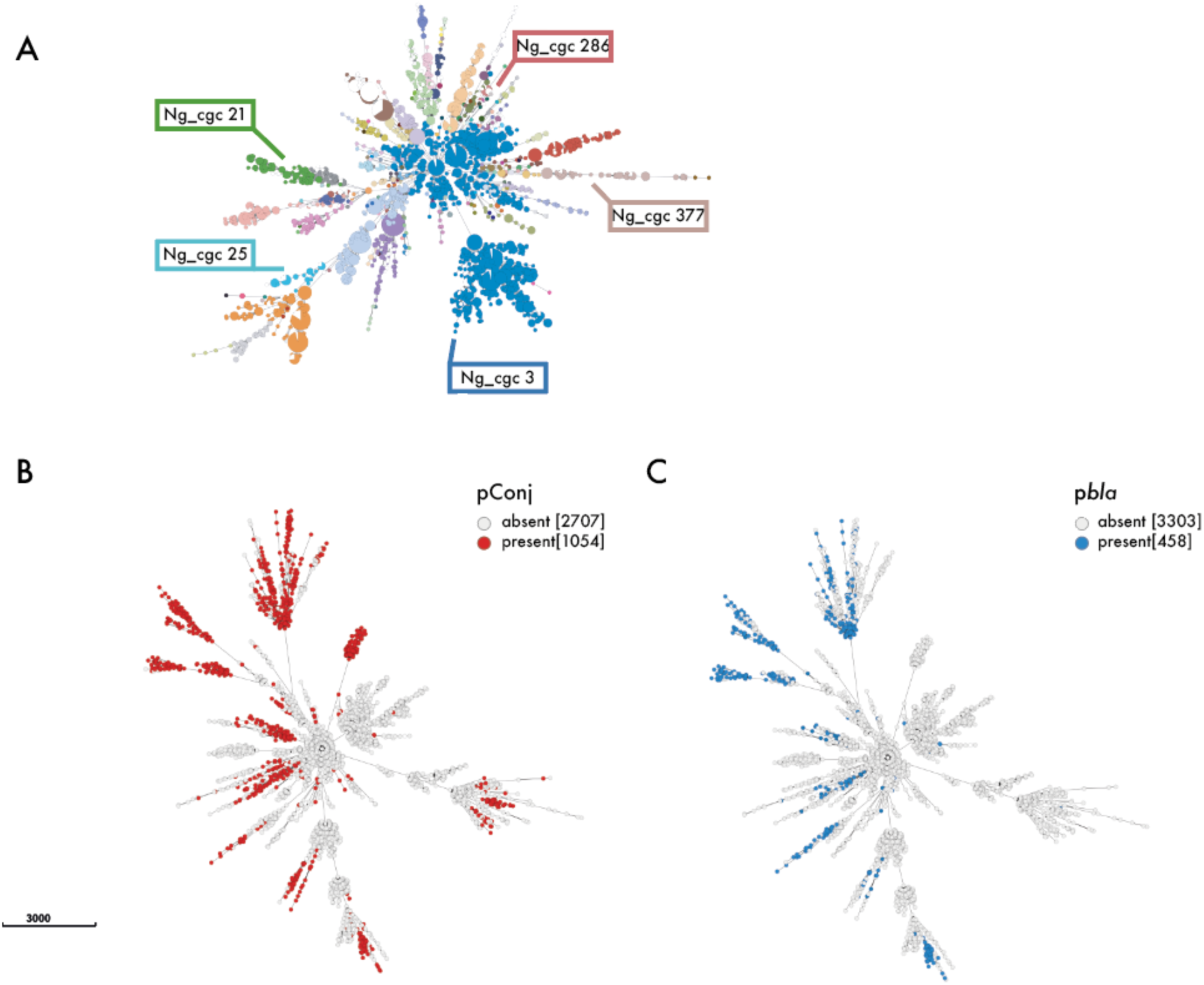
(A-C) Minimum spanning trees of 3,7C1 gonococcal isolates clustered by the 33 RMS loci (A) and 1,CC8 core genome loci (B, C). Isolates are coloured according to their Ng_cgc_400_ (A), with individual Ng_cgcs, or carriage of pConj (B) and pbla (C) highlighted; the number of isolates carrying pConj and pbla is given in brackets.

We assessed the prevalence of pConj and p*bla* in this dataset; p*bla* was present in 440 isolates (11.7%) with pConj found in 1,032 isolates (27.5%), and were prevalent in certain lineages but not others (**Figure 1B and C**) as described previously^22,23^.

Besides the presence/absence of RMSs, phase variation can affect RMS activity^30,32^ and potentially influence plasmid spread. Therefore, for all RMS loci, we identified features such as the homo or polynucleotide tracts that can mediate phase variation^30^, and the presence of premature stop codons; a total of 15 RMS loci had nucleotide sequence features which could affect their activity/undergo phase variation (**Supplementary Table 4**).

To predict which RMSs affect the spread of plasmids, we examined the correlation between variation in RMS loci, including phase-variable features, and the presence/absence of p*bla* and pConj using logistic regression, which is suited to predicting binary outcomes^50^. Each RMS feature (*e.g.* presence of an REase with a premature stop codon) was incorporated as an individual predictor, and logistic regression estimated a regression coefficient for each feature, quantifying the extent to which it contributes to the likelihood of the plasmid being present^50^. A positive regression coefficient indicates a feature is associated with a plasmid, while a negative coefficient reflects association with the lack of a plasmid.

Results of the logistic regression analysis demonstrated that features of i) the NlaIV REase (NEIS1181) and MTase (NEIS1180), ii) the NgoAII REase (NEIS2765), and iii) the HsdS of NgoAV (NEIS2362) were associated with the presence of plasmids (**Supplementary Figure 1A and B, Supplementary Table S5 and S6**). These RMS constitute part of the core genome and have phase-variable features/truncated enzyme variants present in a subpopulation of bacteria. Therefore, we examined their impact on the transfer of p*bla* and pConj in *N. gonorrhoeae* in more detail.

### NlaIV prevents plasmid transfer

NlaIV is a type II RMS which recognises the sequence 5’-GGNN<u>C</u>C-3’^51^. NlaIV_E, the NlaIV REase, is a PD–(D/E)XK endonuclease^52,53^ with D74, E87, and K89 essential for its activity^52^. ON:OFF switching of NlaIV_E could occur due to a poly-T tract within the 5’ end of the gene (**Figure 2A**). *nlaIV_E* with a poly-T tract of 9 (± 3) nt. generates a long NlaIV_E of ∼424 a.a.; frameshift mutations in the poly-T tract can generate a ∼75 a.a. protein, which is predicted to be non-functional as it lacks E87 and K89 (**Figure 2A, B**). *nlaIV_M* has a poly-A tract starting at nt. 793 (Figure 4A); NlaIV_M (423 a.a.) is expected to be functional with poly-A tracts of 9 (± 3) nt.; a frameshift mutation generates short 273 or 291 a.a. versions. Furthermore, a GC insertion at nt. 231 results in a premature stop codon at nt. 271. Linear regression indicated that pConj and p*bla* are more likely to occur in isolates with an inactive, short NlaIV REase (OR = 3.7, chi^2^-test p <0.001 for pConj and OR = 4.8, chi^2^-test p <0.001 for p*bla*), suggesting this RMS impairs plasmid acquisition. Additionally, a premature stop in *nlaIV_M* (NEIS1180) was associated with plasmid presence (regression coefficient = 0.23, log_10_(p) = 18.03 for pConj; regression coefficient = 0.72, log_10_(p) = 38.35 for p*bla*).

**Figure 2:**
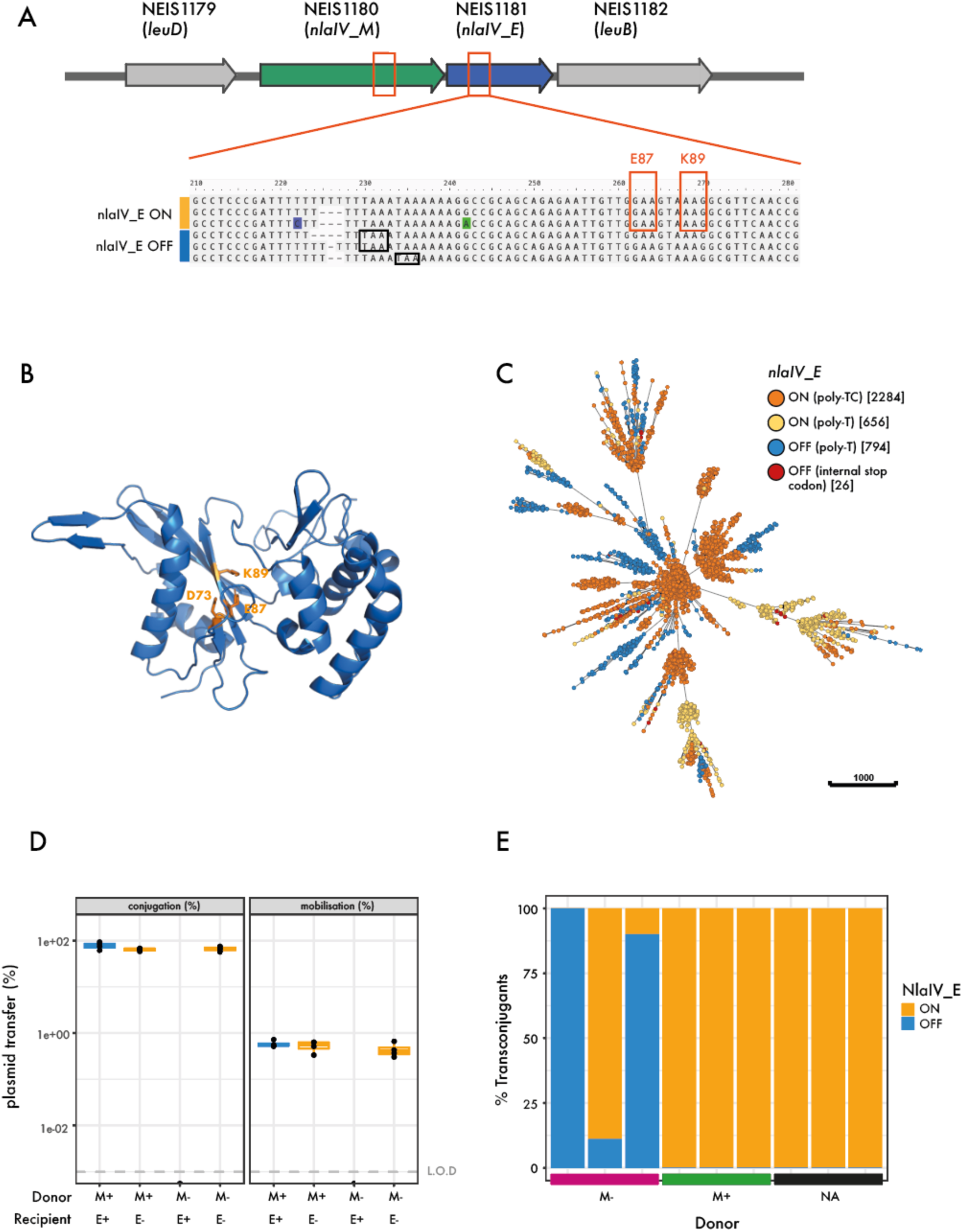
(A) Gene organisation of the nlaIV locus, where nlaIV_M (NEIS1180) and nlaIV_E (NEIS1181) are ffanked by leuD (NEIS117S) and leuB (NEIS1182). Locations of homopolymeric tracts within nlaIV_M and nlaIV_E are indicated with orange boxes. Shown below are the three types of polyT tract in nlaIV_E. Premature stop codons due to changes in the length of the poly-T tract are indicated in black, and codons for the active site residues E87 and K8S are highlighted in orange boxes. (B) AlphaFold3 prediction of NlaIV_E (pTM = 0.SC) with active site residues D73, E87 and K8S indicated in orange. (C) Minimum spanning tree depicting the distribution of nlaIV_E features in the population. (D) Plasmid transfer rates in isogenic matings between donor strains with the methyltransferase ON (M+) or OFF (M-) and a recipient carrying a fixed ON (E+) or OFF (E-) endonuclease. (E) NlaIV-methylated (M+) and non-methylated (M-) strains (FA10S0 and 445CS, respectively) were used as donors of pConj to strain C5C00, which harbours nlaIV_E with an uninterrupted poly-T tract. nlaIV_E in all transconjugants was sequenced to determine the poly-T tract. Recipient only, without the presence of the donor (NA), was used as a control in this experiment.

We investigated the distribution of *nlaIV_E* variation within the gonococcal population. Despite the poly-T tract in *nlaIV_E*, isolates encoding long or short NlaIV_E cluster separately on the minimum spanning tree (**Figure 2C**), suggesting it is not subject to high-frequency phase variation. Furthermore, in 68.2% of isolates, the poly-T tract is interrupted by a synonymous T/C substitution (5’-TTTCTTTTT-3’, **Figure 2A and 2C**, TTT/TTC encode phenylalanine), which maintains the reading frame and likely eliminates phase variation.

To define the ability of NlaIV to hinder plasmid transfer, we constructed isogenic *N. gonorrhoeae* FA1090 strains with active/inactive NlaIV_M (*nlaIV_M*_ON_/*nlaIV_M*_OFF_) and NlaIV_E (*nlaIV_E*_ON_/*nlaIV_E*_OFF_). For the *nlaIV_E*_OFF_ strain, an additional T was introduced at the 3ʹ end of the interrupted poly-T tract (generating 5’-TTTTCTTTTT-3’) to generate a short NlaIV_E (71 a.a.). To limit phase variation of NlaIV_M, the poly-A tract was interrupted by a synonymous A/G substitution at nt. 6 of the tract (5’-AAAAAGAAA-3’, AAA/AAG encode lysine) in the *nlaIV_M*_ON_ strain; an A was removed from the end of the tract (5’-AAAAAGAA-3’) to generate the *nlaIV_M*_OFF_ strain.

Matings were performed between pConj.1 and p*bla*.1 (the commonest variants^27,28^) containing donors with either an active or inactive NlaIV MTase and an inactive REase, and recipients with the MTase ON and the REase either ON or OFF (**Figure 2D**). Donors contained *pilD::ermC,* while recipients harboured *pilD::aph(3)* to select donors and recipients/transconjugants, and prevent plasmid transfer by transformation^54^. In all matings with an *nlaIV_M*_ON_ donor, there was no difference in plasmid transfer into an *nlaIV_E*_ON_ or *nlaIV_E*_OFF_ recipient. Strikingly, there was no detectable transfer of p*bla* and pConj from a donor lacking NlaIV methylation into recipients with an active NlaIV REase (<0.0001%, **Figure 2D**). In contrast, the transfer of plasmids from the NlaIV methylation-defective donor was unimpeded into a recipient with inactive REase, demonstrating that active NlaIV_E prevents uptake of unmethylated p*bla* and pConj, consistent with logistic regression.

We reasoned that phase variation of *nlaIV_E* could create subpopulations of bacteria that could acquire plasmids from a donor lacking NlaIV methylation. To test this, we characterised transconjugants arising from matings between a recipient carrying an active yet phase-variable *nlaIV_E* (*i.e.* isolate *N. gonorrhoeae* 65600) with a donor with either *nlaIV_M*_ON_ or *nlaIV_M*_OFF_ (isolate 44569, which has *nlaIV_E*_ON_); the *nlaIV* locus was amplified and sequenced in transconjugants. Of note, transconjugants from the mating between a recipient with an active MTase all have the poly-T tract of *nlaIV_E* ON as in the parental strain. In contrast, 68% of transconjugants from the matings with the unmethylated donor had *nlaIV_E* switched OFF (**Figure 2E**), highlighting that phase variation can allow uptake of non-NlaIV methylated plasmids.

Even though the NlaIV RMS has a profound effect on plasmid transfer, only 49 of 3,761 isolates (1.3%) are predicted to have a truncated MTase. Therefore, the overwhelming majority of *N. gonorrhoeae* population is methylated by NlaIV_M, so this RMS is unlikely to be a significant barrier to intraspecies plasmid flow, but instead might have a profound effect at preventing plasmids from being acquired from species lacking this RMS. Of note, NlaIV is found in *N. meningitidis*, the likely source of pConj for the gonococcus^24^.

### NgoAII is unlikely to impede plasmid transfer between gonococci

The type II RMS NgoAII recognises the sequence 5’-GG<u>C</u>C-3’^30^. The NgoAII REase, NgoAII_E (NEIS2765), has been classified as PD-(D/E)XK endonuclease^55^. NgoAII_E can either be full-length (278 a.a., NgoAII_E_278_) or truncated (134 a.a., NgoAII_E_134_) due to insertion of an A at nt. 388 (**Figure 3A**). The MTase NgoAII_M is encoded downstream of *ngoAII_E* by NEIS2722.

**Figure 3:**
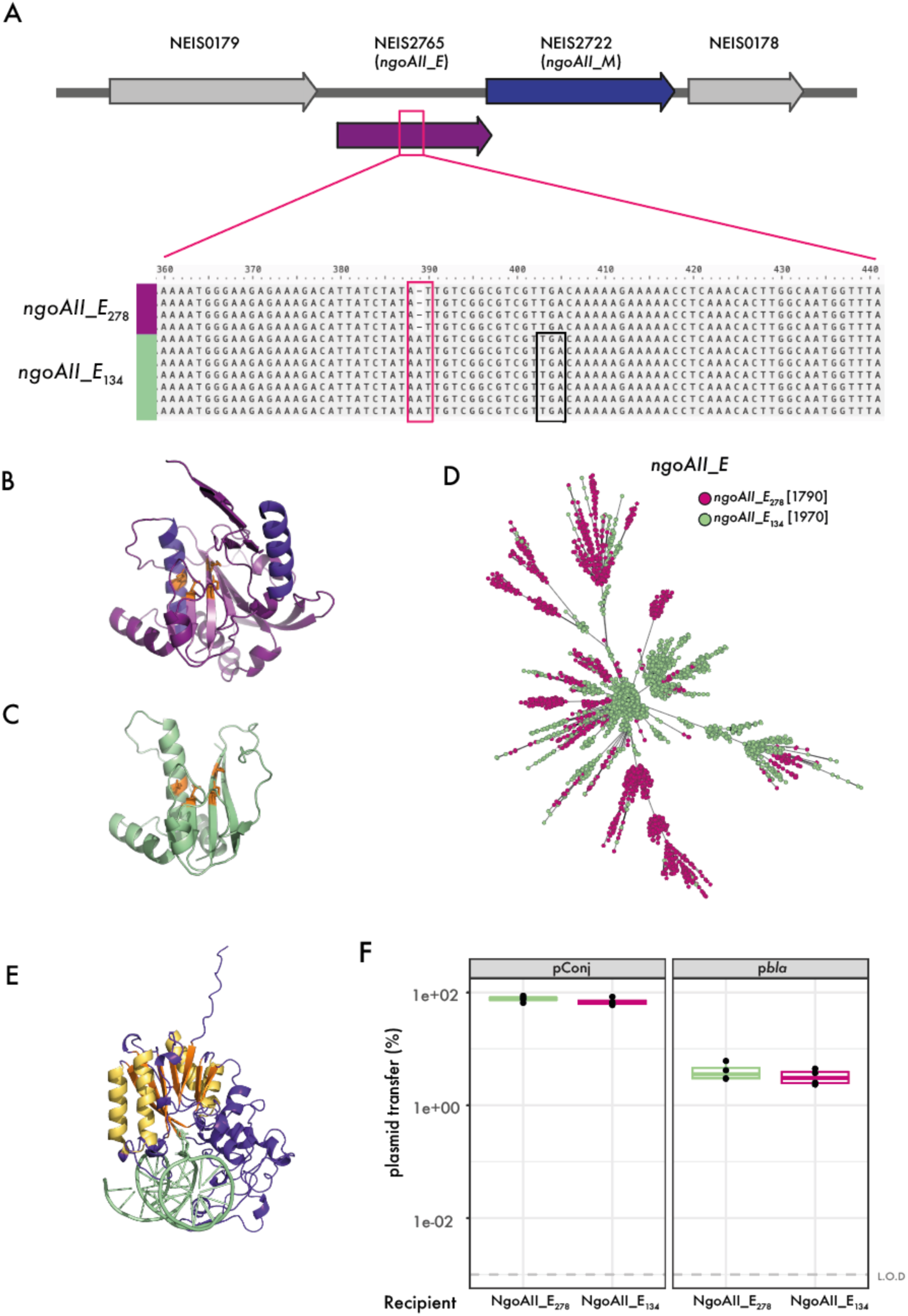
(A) Genetic organisation of the ngoAII locus. NEIS27C5 (ngoAII_E) and NEIS2722 (ngoAII_M) are ffanked by NEIS017S, encoding a putative inner membrane protein, and NEIS0178, encoding a ribosome recycling factor. An adenine insertion at position 388 (pink box) results in a frameshift mutation and a premature stop codon (highlighted in black). (B, C) AlphaFold3 predictions of NgoaAII_E278 (B) and NgoAII_E134 (C). The secondary structure elements of the conserved core fold of PD-(D/E)XK nucleases are shown in purple (α-helices) and violet (β-sheet), and the key active site residues E3C, D73, E83, K85 are shown in orange. (D) Minimum spanning tree of 3,7C1 isolates that were clustered according to core genomic allelic differences (cgMLST v1). Individual isolates are represented as dots that are coloured according to the ngoAII_M allele carried. (E) AlphaFold3 prediction of NgoAII_M with GGCC motif-containing dsDNA. The first C of the GGCC motif is flipped into the active site of the methylase. Secondary structure elements conserved between class I MTases are indicated in yellow (α-helices) and orange (β-sheet). (F) Plasmid transfer into isogenic recipients that either harbour full-length or truncated NEIS275. Wild-type FA10S0, which carries NEIS2722 allele 1, served as the donor. There was no significant difference in pConj or pbla transfer into the different donors (Welch two-sample t-test p = 0.28 and 0.41, respectively).

**Figure 4:**
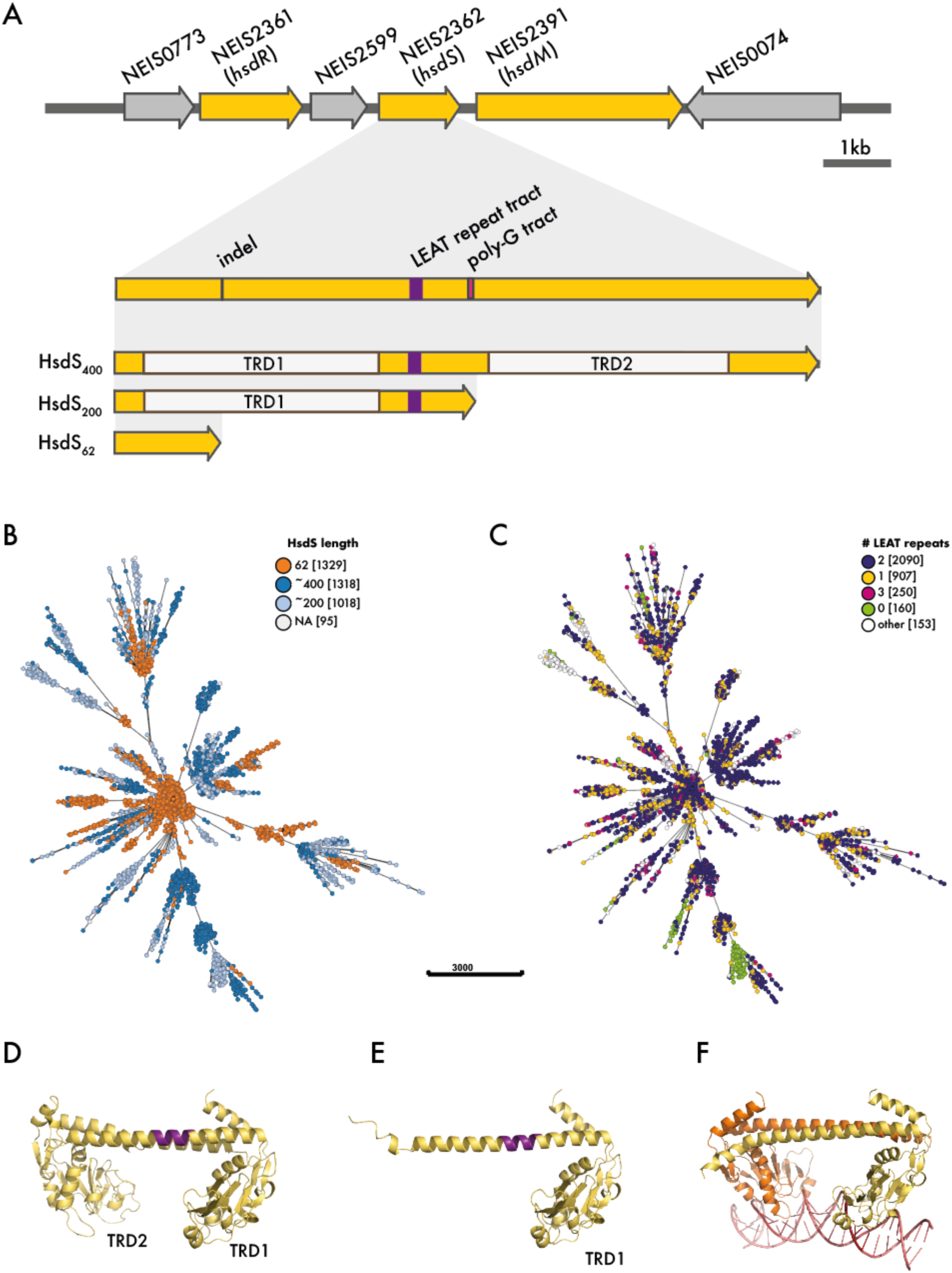
(A) Gene organisation of the ngoAV locus, where hsdM (NEIS23C1), hsdS (NEIS23C2) and hsdR (NEIS23S1) are ffanked by the genes encoding an ADP-D-beta-D heptose epimerase (NEIS0073) and a putative ATP-dependent protease (NEIS0074). NEIS25SS encodes the DNA damage-inducible protein D. Shown below are the three variable regions within hsdS resulting in HsdS400, HsdS200 and HsdSC2: an indel at position 18C (leading to an inactive enzyme), a region with repeats of the nucleotide sequence encoding the amino acids ’LEAT’ (purple rectangle) and a poly-T tract (pink rectangle). (B,C) Minimum spanning trees of 3,7C1 gonococcal isolates displaying HsdS length in amino acids (B) and the number of LEAT repeats (C) in the population. Incomplete hsdS loci, where the number of LEAT repeats and HsdS length could not be determined, are indicated as NA. (D, E) AlphaFold3 predictions of HsdS400 (F) and HsdS200 (F). LEAT repeat regions are indicated in purple. (G) Structure prediction of HsdS200 dimer with dsDNA containing the recognition sequence GCAN8TGC.

We generated structural models of NgoAII_E_278_ and NgoAII_E_134_ with AlphaFold3. NgoAII_E_278_ shows the conserved folds of a PD-(D/E)XK endonuclease with a β-sheet flanked by two α-helices and all predicted active site residues (E36, D73, E83, K85, **Figure 3B**)^55,56^. While NgoAII_E_134_ retains all active site residues (**Figure 3C**), part of its second α-helix is lost. Most PD–(D/E)XK REases act as dimers^57^, with AlphaFold3 predicting that NgoAII_E_278_ dimerisation is facilitated by its C-terminal domain (**Supplementary Figure 2A**). However, NgoAII_E_134_ lacks the dimerisation region (**Figure 3C**), so is likely to be inactive.

Surprisingly, logistic regression indicated that the presence of the full-length active NgoAII REase was associated with pConj and p*bla* carriage (regression coefficients 1.31, log_10_(p) = 70.52 and 0.96, log_10_(p) = 36.41, respectively, **Supplementary Figure 1).**

NgoAII_E_278_ and NgoAII_E_134_ are present at approximately equal frequency in the gonococcal population (47.8% and 52.2% of isolates, respectively, **Figure 3D**) and isolates with the same NgoAII_E cluster together in the population, indicating that it is unlikely to be phase-variable. Additionally, NgoAII_M is highly conserved in the gonococcus; alleles 1, 4, 7, 8, 11, 12 and 31 have synonymous mutations, and account for 77.8% of sequences (**Supplementary Figure 2B**); allele 3 in 15.7% of isolates encodes for the MTase with a single G145R substitution. Using AlphaFold3, NgoAII_M is predicted to possess the conserved αβα motif of Class I MTases, consisting of a seven β-sheets flanked by α-helices (**Figure 3E**)^58^. Modelling of NgoAII_M with dsDNA containing its 5ʹ-GGCC-3ʹ motif reveals an electropositive groove predicted to interact with dsDNA and flipping of the target C within the motif (**Figure 3E, Supplementary Figure 2C**). The G145R substitution is located on the surface of the enzyme, distant from the DNA-binding groove/active site (**Supplementary Figure 2D**), so is unlikely to affect enzyme activity. Consequently, 99.9% of isolates in the dataset are predicted to encode an active MTase, suggesting that NgoAII does not limit plasmid transfer between gonococci.

We constructed isogenic FA1090 strains with *ngoAII_E*_278_ and *ngoAII_E*_134_. Matings were performed between FA1090 *pilD::ermC* pConj.1 and p*bla*.1, with NgoAII_M allele 1 as the donor, and FA1090 *pilD::aph(3)* with *ngoAII_E*_278_ or *ngoAII_E*_134_ as the recipient. As expected from a donor with an active methyltransferase, there was no difference in pConj and p*bla* transfer rates into FA1090 expressing full-length NgoAII_E or the truncated enzyme (**Figure 3F**, Welch two-sample t-test p = 0.28 and 0.41, respectively), consistent with NgoAII having little or no impact on plasmid spread in *N. gonorrhoeae*.

### Variation in NgoAV hsdS results in distinct methylation patterns

Variation in NgoAV *hsdS* arises through i) a region encoding different numbers of the amino acids, LEAT^59^, ii) a poly-G tract (of 6 to 15 nt.) starting at ∼ nt. 612 of *hsdS,* which generates a long (HsdS_400_) or short (HsdS_200_) protein^32^, and iii) insertion of a T at nt. 186, resulting in HsdS of 62 a.a. (HsdS_62_, **Figure 4A**). The activity of NgoAV is altered by *hsdS* variation^30,32^. HsdS_200_ methylates the palindromic sequence, 5ʹ-GC<u>A</u>(N_8_)TGC-3ʹ, HsdS_62_ is inactive, while changes in the number of LEAT repeats alter spacing of the non-palindromic motif, 5ʹ-GC<u>A</u>N_X_GTC-3ʹ/5ʹ-G<u>A</u>CN_X_TGC-3ʹ^32,60^.

Logistic regression indicated that HsdS without LEAT is associated with pConj presence (regression coefficient = 0.86, log_10_(p) = 23.98), but p*bla* absence (regression coefficient = -0.29, log_10_(p) = 31.51, **Supplementary Figure 1A and B**). Initially, we investigated the distribution of *hsdS* variants in the 3,761 gonococcal isolates. Sequences encoding HsdS_62_, HsdS_200_ and HsdS_400_ were present at approximately equal frequency (35.3%, 27.2% and 35.0%, respectively, **Figure 4B**). Of note, isolates that cluster together on the tree tend to encode the same length HsdS (**Figure 4B**), suggesting that neither the T insertion at nt. 186 nor the poly-G tract undergo significant phase variation. In contrast, the number of LEAT repeats in HsdS varies between closely related isolates, indicating that this feature is phase-variable (**Figure 4C**); as expected due to their lack of repeated sequence, exceptions are isolates expressing HsdS with 0 or 1 LEAT (**Figure 4C**).

The most frequent variants were HsdS_400_/2LEAT (23.6% of isolates, **Table 1**), followed by HsdS_200_/2LEAT (13.5%) and HsdS_400_/1LEAT (10.0%) (**Figure 4C**, **Table 1**). HsdS/0LEAT was HsdS_200_ in 96.9% of sequences with 0 LEAT. To understand how these changes in HsdS affect methylation, we generated isogenic strains of *N. gonorrhoeae* FA1090 with these four prevalent HsdS variants. Strains were sequenced with Nanopore, and data were processed with Dorado and Modkit to detect chromosomal and plasmid m6A methylation (**Supplementary Table 7**). All strains showed evidence of methylation by NgoAXVI (5ʹ-GGTG<u>A</u>-3ʹ) and NgoAXII (5ʹ-GCAG<u>A</u>-3ʹ)^30^, which are predicted to be active in FA1090. Strains encoding HsdS_400_ also showed methylation of 5ʹ-GC<u>A</u>N_X_GTC-3ʹ/5ʹ-G<u>A</u>CN_X_TGC-3ʹ, with >99.5% of motifs in the chromosome/plasmids modified. Of note, the spacer length between methylated sites was altered by the number of LEATs, with N = 6 nts. with one LEAT, and = 7 nts. with two LEATs. In contrast, HsdS_200_/2LEAT methylates a palindromic sequence, 5ʹ-GC<u>A</u>N_8_TGC-3ʹ (99.3% of motifs in the chromosome/plasmids modified, **Supplementary Table S7, Supplementary Figure 3**)^30^.

**Table 1:**
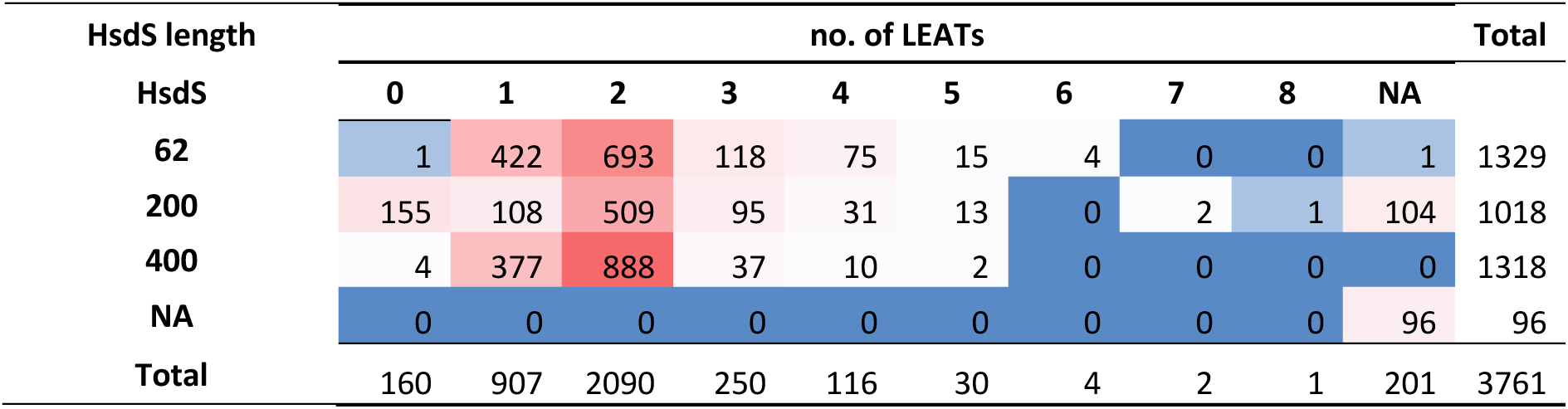
Prevalence of different NgoAV HsdS features (length of the encoded HsdS, and number of repeats of the amino acid sequence ’LEAT’ within HsdS) encoded by isolates in the dataset. Prevalent combinations are highlighted in red, while infrequent combinations are shown in blue. NA indicates incomplete hsdS loci where features could not be determined.

| HsdS length | no. of LEATs |  |  |  |  |  |  |  |  |  | Total |
| --- | --- | --- | --- | --- | --- | --- | --- | --- | --- | --- | --- |
| HsdS | 0 | 1 | 2 | 3 | 4 | 5 | 6 | 7 | 8 | NA |  |
| 62 | 1 | 422 | 693 | 118 | 75 | 15 | 4 | 0 | 0 | 1 | 1329 |
| 200 | 155 | 108 | 509 | 95 | 31 | 13 | 0 | 2 | 1 | 104 | 1018 |
| 400 | 4 | 377 | 888 | 37 | 10 | 2 | 0 | 0 | 0 | 0 | 1318 |
| NA | 0 | 0 | 0 | 0 | 0 | 0 | 0 | 0 | 0 | 96 | 96 |
| Total | 160 | 907 | 2090 | 250 | 116 | 30 | 4 | 2 | 1 | 201 | 3761 |

To understand how *hsdS* variation affects its activity, we predicted structures of HsdS_400_ and HsdS_200_ using AlphaFold3^61^ (**Figure 4D and E**, pTM = 0.66 and pTM = 0.77, respectively). Similar to the structure of HsdS from the type I RMS EcoR124I^10^, HsdS_400_ is predicted to have two target recognition domains (TRD1 and TRD2, a.a. 18 to 153 and 219 to 356, respectively, **Figure 4D**)^32^, which share a high degree of structural similarity (RMSD, 1.736 Å) and are separated by two α-helices (**Figure 4D**). The presence of different numbers of LEAT repeats which occur in an α-helix is predicted to change the relative spacing between and pitch of the TRDs (**Supplementary Figure 4**). For example, spacing between the TRDs in predicted structures of HsdS_400_ with 1 and 2 LEAT repeats was 72.8 Å and 76.5 Å, respectively (measured from S154 of TRD1 to N210/214/218 of TRD2). This offers a structural explanation for the change in motif spacing with an alteration in LEAT repeats, with HsdS_400_ acting as a molecular ruler extended and retracted by alterations in the number of LEATs.

Even though HsdS_200_/2LEAT only possesses a single TRD1 and one α-helix (**Figure 4E**), it still methylates a palindromic sequence, 5ʹ-GC<u>A</u>N_8_TGC-3ʹ. As 5ʹ-GC<u>A</u>-3ʹ is also methylated by HsdS_400_, TRD1 likely recognises this sequence, while methylation of palindromic sequences by HsdS_200_/2LEAT suggests that it dimerises. This is supported by AlphaFold3 predictions of HsdS_200_ with dsDNA containing the 5ʹ-GCAN_8_TGC-3ʹ motif. Dimerisation between two HsdS_200_ occurs along the length of the α-helix (**Figure 4F**, ipTM = 0.55), resulting in a structure with similar overall architecture as HsdS_400_ (executive RMSD 2.322 Å). This generates an atypical M_2_S_2_ specificity complex with two TRD1s and hence a palindromic motif. NgoAV methylation was not detected in the strain expressing HsdS_200_/0LEAT, indicating that this variant is inactive, and dimerisation depends on LEATs in the α-helix.

### NgoAV has a minimal impact on plasmid transfer

As HsdS variation is associated with plasmid presence/absence and alters methylation patterns, we next examined its effect on plasmid transfer. Matings were performed between isogenic *N. gonorrhoeae* FA1090 donors and recipients expressing different versions of HsdS (*i.e.* HsdS_400_/2LEAT, Hsd_400_/1LEAT, HsdS_200_/2LEAT, HsdS_200_/0LEAT, **Figure 5**). Donors contained *pilD::ermC,* p*bla*.1, and pConj.1, while recipients harboured *pilD::aph(3)*. Results demonstrate that plasmid transfer differed between matings, with significant differences between both donors and recipients and a significant interaction between donor and recipient (two-way factorial ANOVA including the fixed effects of donor and recipient and their interaction, p < 0.001 for donor, recipient and their interaction for both p*bla* and pConj). However, conjugation rates were reduced by less than 50% compared to matings between donors and recipients with matched *hsdS*. p*bla* mobilisation was not dramatically affected by differences in *hsdS* sequences, consistent with their being few NgoAV recognition sites on this plasmid (**Supplementary Figure 3**).

**Figure 5:**
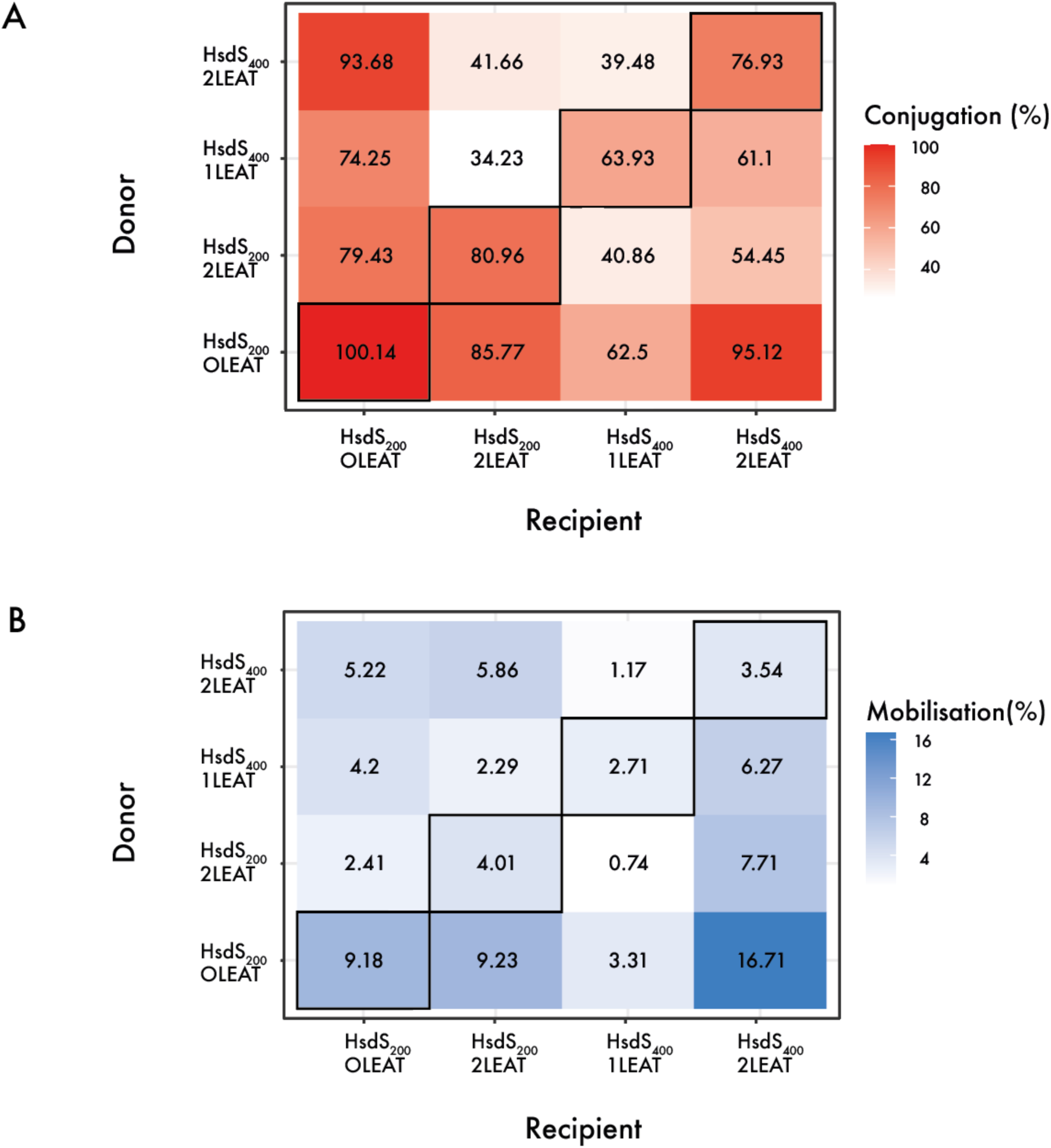
Heatmaps displaying the transfer rates of pConj (A) and pbla (B) in matings between isogenic FA10S0 strains with different hsdS. Darker colours indicate higher plasmid transfer, and numbers in boxes give the average of matings performed on three occasions. Isogenic matings along the diagonal are highlighted.

Consistent with our findings that HsdS with 0 LEAT is both inactive and associated with pConj carriage (**Supplementary Figure 1A**), the strain expressing HsdS_200_/0LEAT (which accounts for >95% of isolates with 0 LEAT) is the most effective recipient of pConj and p*bla* (Tukey multiple comparison of means, p < 0.01 for pConj transfer compared with HsdS_200_/2LEAT and HsdS_400_/1LEAT, p = 0.06 for the HsdS_400_/2LEAT recipient, and p < 0.001 for p*bla* transfer); plasmid transfer into the strain expressing HsdS_200_/0LEAT often exceeded those observed between strains with matched *hsdS* alleles. However, counterintuitively, this strain was also an efficient donor (**Figure 5**, Tukey multiple comparisons of means, p < 0.001 for all pairwise comparisons with unmatched *hsdS* alleles and including HsdS200/0LEAT, for both p*bla* and pConj).

In summary, HsdS variation influences plasmid transfer, with the number of LEAT repeats extending/retracting the HsdS molecular ruler, while HsdS can switch NgoAV recognition from a non-palindromic to a palindromic motif by assuming a unique HsdM_2_S_2_ stoichiometry.

### Plasmid transfer is affected by factors other than RMSs

The combinatorial effects of core, non-phase-variable RMSs in the gonococcus (**Figure 1A**) could limit the spread of the plasmids between different lineages. To investigate this, we assessed the transfer of pConj and p*bla* between clinical isolates from Ng_cgc_400_ lineages 3, 21, 25, 286 and 377 (**Supplementary Figure 5**). When clustering our dataset according to their RMS profiles, these Ng_cgc_400_s form distinct clusters, indicating they possess distinct alleles of their RMSs (**Figure 1A**). Furthermore, these lineages differ in their plasmid prevalence: Ng_cgc_400_s 21, 25 have high and Ng_cgc_400_ 3, 286 have low plasmid carriage, while Ng_cgc_400_ 377 is unusual as it has a high p*bla* (52.8%, 140/265) but low pConj (1.9%, 5/265) carriage (**Supplementary Table S8**).

For each lineage, a plasmid-free isolate was selected, and a donor generated by introducing p*bla*.1 and pConj.1, as well as *pilD::ermC*. Recipients were constructed by replacing *pilD* with *aph(3).* In line with the diversity of NgoAII and NlaIV, all of the isolates had active NlaIV and NgoAII methylases; *nlaIV_E* was off in one isolate (NG149) and three isolates (NG015, NG149 and FA1090) encoded truncated NgoAII_E (**Table 2**, full RMS allelic profiles are given in **Supplementary Table S9**). All isolates expressed NgoAV HsdS_200_/2LEAT except for NG015, which encoded HsdS_400_/2LEAT.

**Table 2:** NgoAII, NgoAV and NlaIV features of isolates used for the mixed isolate matings.

| Isolate | Ng_cgc<br>400 | NgoAV HsdS | NlaIV_M | NlaIV_E | NgoAll_E | NgoAll_M |
| --- | --- | --- | --- | --- | --- | --- |
| NG015 | 3 | HsdS<sub>400</sub>/2LEAT | + | + | - | + |
| NG114 | 25 | HsdS<sub>200</sub>/2LEAT | + | + | + | + |
| NG149 | 377 | HsdS<sub>200</sub>/2LEAT | + | - | - | + |
| 2086_K | 21 | HsdS<sub>200</sub>/2LEAT | + | + | + | + |
| FA1090 | 286 | HsdS<sub>200</sub>/2LEAT | + | + | - | + |

We performed matings between all combinations of isolates measuring pConj and p*bla* transfer (**Figure 6A/B**). Comparison of isogenic matings demonstrated that the rate of plasmid transfer was highly variable. FA1090, NG149 and 2086_K gave high pConj and p*bla* transfer (>69% and >2.9%, respectively), while plasmid transfer for NG015 or NG114 was an order of magnitude lower (conjugation rates <7.4% and mobilisation rates <0.38%), indicating that features other than RMSs affect plasmid transfer within gonococcus.

**Figure 6:**
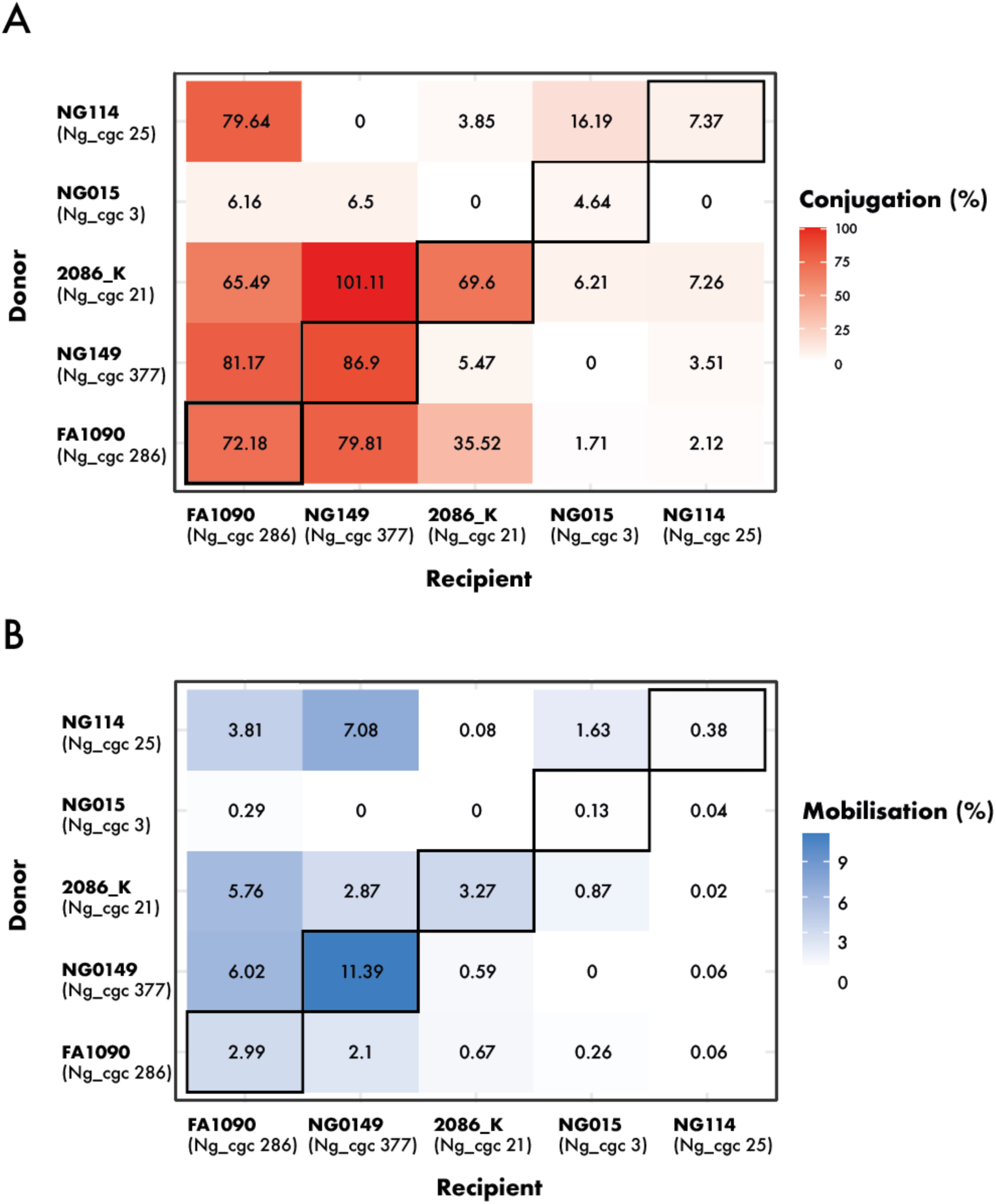
(A, B) Heatmaps of the pConj conjugation (A) and pbla mobilisation frequency (B) between different donors and recipients. Isogenic matings along the diagonal are highlighted. Darker colours indicate higher plasmid transfer, and numbers in boxes give the average of matings performed on three occasions.

Several non-isogenic matings had equivalent or higher plasmid transfer rates than isogenic matings. Most notable, pConj and p*bla* transfer from NG114 into FA1090 was 79.6% and 3.8%, respectively, while isogenic NG114 matings gave conjugation and mobilisation frequencies of only 7.4% and 0.38%, with NG114 an inefficient recipient (pConj transfer <7.4%, p*bla* transfer <0.4, **Figure 6A/B**), while FA1090 was an efficient recipient (**Figure 6A/B**). The highest transfer rates for both pConj and p*bla* were with NG149 as the recipient, as expected, since NG149 is the only strain lacking NlaIV and NgoAII REases (**Table 2**).

Transfer rates also depended on the donor. NG015 was an inefficient donor and was the only strain with the active HsdS_400_/2LEAT (**Figure 5**). In contrast, matings with 2086_K as the donor resulted in comparable or higher plasmid transfer as observed in isogenic matings. Taken together, even with this limited number of isolates, plasmid transfer between gonococcal lineages cannot be explained by host-encoded barriers to HGT alone.

## DISCUSSION

RMSs have been long regarded as a bacterial innate immune system, preventing the acquisition of foreign DNA and phage infection. Here, we examined the role of RMSs on the transfer plasmids in *N. gonorrhoeae*, a bacterium with an exceptionally large number of RMSs compared to other species with similar genome sizes^31^. While in many bacteria, including the meningococcus^16^, the repertoire of distinct RMSs dictates HGT between lineages, most *N. gonorrhoeae* RMS loci are part of the core genome, indicating that the repertoire of RMSs in the gonococcus does not limit genetic transfer between lineages. Instead, our analyses reveal that phase variation in core RMSs affects their activity and the flow of the important AMR plasmids, pConj and p*bla*.

The type II RMS NlaIV was highly effective at preventing plasmid transfer from a donor lacking NlaIV methylation. The NlaIV REase is phase-variable due to a homopolymeric tract in the 5’ end of the gene. Matings with a recipient with an uninterrupted homopolymeric tract yielded only rare transconjugants that had undergone ON to OFF switching of the Rease. As such, NlaIV is a potent restriction barrier that effectively discriminates between self and non-self DNA, and limits HGT into the gonococcus. *N. meningitidis*, which carries ancestral variants of pConj, also encodes NlaIV^56^, suggesting that this shared restriction profile permitted pConj transfer from the meningococcus into *N. gonorrhoeae*^24^, and may explain why the gonococcus, despite its natural competence, harbours so few plasmids. Once in *N. gonorrhoeae*, plasmids would be methylated by NlaIV MTase, allowing for their further spread in the gonococcal population. Although *nlaIV_M* also contains a homopolymeric tract, the overwhelming majority of isolates are predicted to express an active MTase. Therefore, while NlaIV could limit the acquisition by *N. gonorrhoeae* of plasmids from other species lacking NlaIV RMS, it is unlikely to limit p*bla* and pConj transfer between gonococci.

Logistic regression analysis also implicated the NgoAII REase with the presence of plasmids, although counterintuitively, a functional endonuclease was associated with plasmid carriage, suggesting this result might be due to genetic linkage to other genetic factors involved in plasmid transfer. Similar to NlaIV, the NgoAII MTase is highly conserved across gonococci, and is expected to be non-functional in only three of 3,761 isolates. Thus, most plasmids circulating in gonococci should be methylated by NgoAII, and the NgoAII REase should not affect plasmid transfer in the gonococcal population, consistent with our plasmid transfer results.

Three regions in NgoAV *hsdS* generate remarkable diversity in its activity through distinct mechanisms. First, there are HsdSs of different lengths; while HsdS_62_ is inactive, both longer versions can be active. Second, HsdS_200_/2LEAT is predicted to dimerise to reconstitute a functional subunit with two TRD1s and a similar architecture to HsdS_400_; the palindromic sequence modified by HsdS_200_/2LEAT supports the notion that there are two TRD1s in a non-canonical HsdM_2_S_2_ MTase. Furthermore, the LEAT repeat acts as a molecular ruler that regulates the atomic distance between TRDs, and therefore the spacing of NgoAV’s motifs. Of note, we did not detect NgoAV methylation in HsdS/0LEAT strains, indicating that the absence of a LEAT sequence inactivates the RMS. The HsdS ruler coupled its reversible dimerisation establishes an unusual example of phase variation that generates structural diversification and changes in motif symmetry and spacing rather than ON:OFF switching.

Using engineered *N. gonorrhoeae* strains, variation in NgoAV HsdS had a detectable effect on plasmid transfer, albeit less marked than NlaIV. However, the effect of NgoAV on plasmid transfer might have more significant effects during gonococcal infection with different bacterial density, growth and selective pressures. The strain with NgoAV inactive (*i.e.* HsdS200/0LEAT) was an efficient recipient, independent of donor, as predicted by logistic regression analysis. Conversely, this strain also proved to be an effective donor. The activity of an RMS with an overlapping motif could explain the high transfer rates of the donor with an inactive HsdS. For instance, NgoAVII modifies 5’-GACN_5_TGA-3’^8^ which includes the sequence recognised by NgoAV HsdS TRD1. However, these was no detectable NgoAVII methylation in FA1090, indicating that the RMS is inactive in this strain. Certain gonococcal RMSs mediate epigenetic gene regulation^33,34,62^, and NgoAV methylation has a genome-wide effect on transcription^33^. Therefore, expression of plasmid-encoded genes involved in conjugation might differ between strains with distinct *hsdS* alleles, affecting conjugation rates.

*N. gonorrhoeae* displays a non-clonal population structure due to frequent intraspecies HGT^63–65^. The multitude of RMSs, including NlaIV, in gonococci should, on the other hand, provide a potent barrier against interspecies HGT, and might explain why the gonococcus has such a limited repertoire of plasmids, while other high priority AMR pathogens harbour a plethora of mobile genetic elements^66–68^.

As the population structure of the gonococcus can confound correlations detected by logistic regression, we directly tested the predicted the effect of RMSs with isogenic and non-isogenic matings. The patterns of plasmid transfer between clinical isolates could not be simply explained by host-encoded barriers to HGT, as matings between isogenic pairs did not consistently give the highest transfer rates. Instead, other factors influence the activity of plasmid transfer. This could be mediated by transcriptional regulation of the plasmid, or maybe an inherent property of the bacterial surface affecting HGT. Thus, other currently uncharacterised factors likely influence the transfer of plasmids between gonococci, and contribute to their uneven distribution in the population.

In summary, we identified 33 RMS loci in our dataset of 3,761 isolates, with most of them forming part of the core gonococcal genome. The conserved yet extensive repertoire of RMSs in *N. gonorrhoeae* shapes inter and intra-species plasmid dynamics and likely constitutes an effective barrier against the acquisition of plasmids from other species. Exceptions will occur when other species harbour homologous RMSs, for instance NlaIV in other *Neisseria* spp. ^51,69,70^. Additionally, the interplay between different phase variation events within NgoAV HsdS alters spacing, sequence, and symmetry of epigenetic modification. Mechanistically, this occurs through the LEAT repeats affecting the length of a molecular ruler embedded in the α-helices of HsdS, while dimerisation of a truncated version of HsdS leads to modification of palindromic, rather than asymmetric, motifs. This capacity for reversible epigenetic reprogramming provides a link between molecular structure, restriction, and the evolution and spread of resistance.

## Supplementary Figures

**Supplementary Figure 1:**
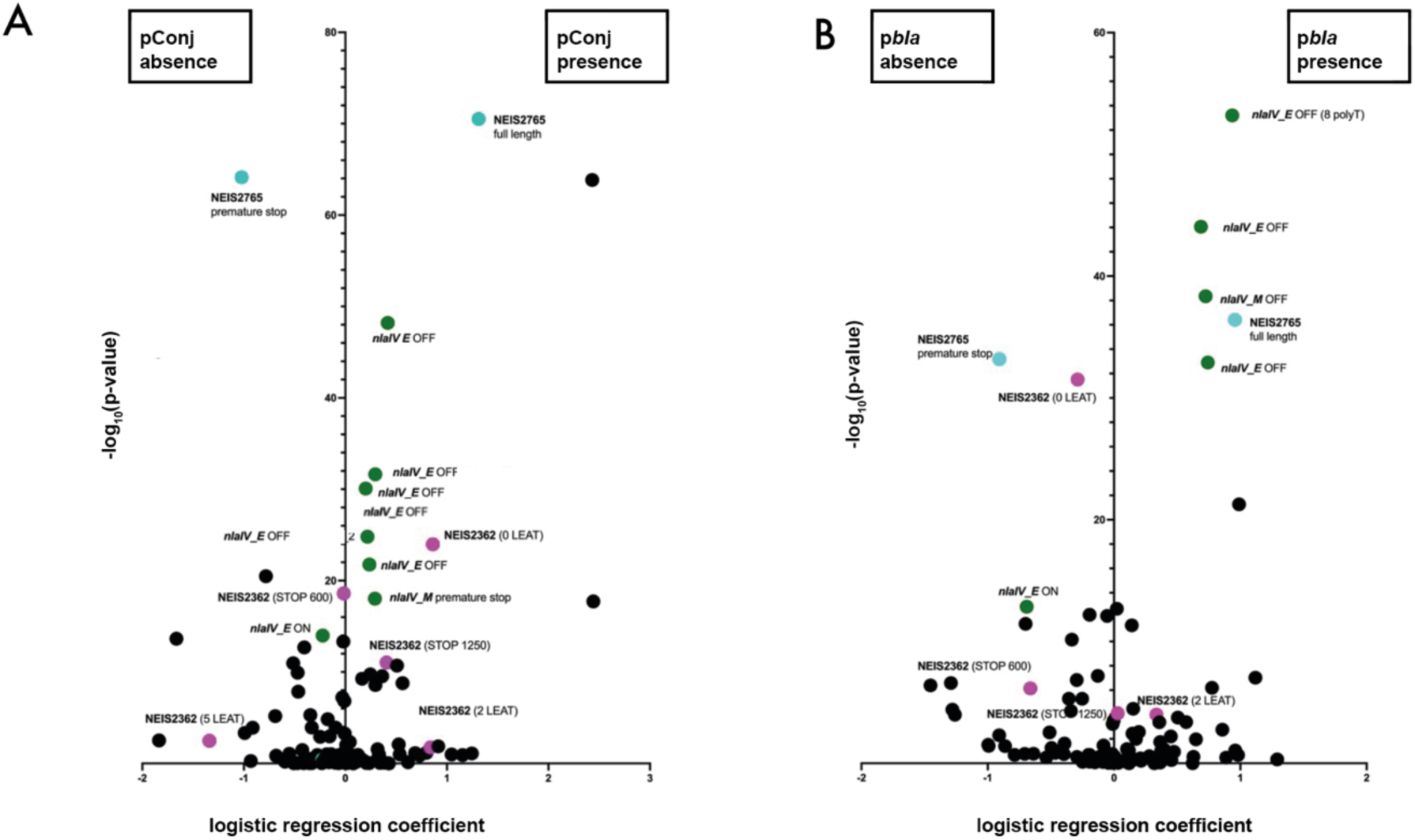
Logistic regression of the correlation of individual features of RMS genes with plasmid presence (positive logistic regression coefficient) or absence (negative logistic regression coefficient). The y-axis indicates the negative logarithm of the p-value of the respective prediction, i.e. the higher the value, the more confident the prediction. Dots represent individual features of RMSs. Features of NEIS2765 (cyan), nlaIV_E and nlaIV_M (green) and NEIS2362 (magenta) are indicated for pConj (A) and pbla (B). Different poly-T tract lengths within nlaIV_E result in inactive NlaIV_E and are indicated as separate datapoints

**Supplementary figure 2:**
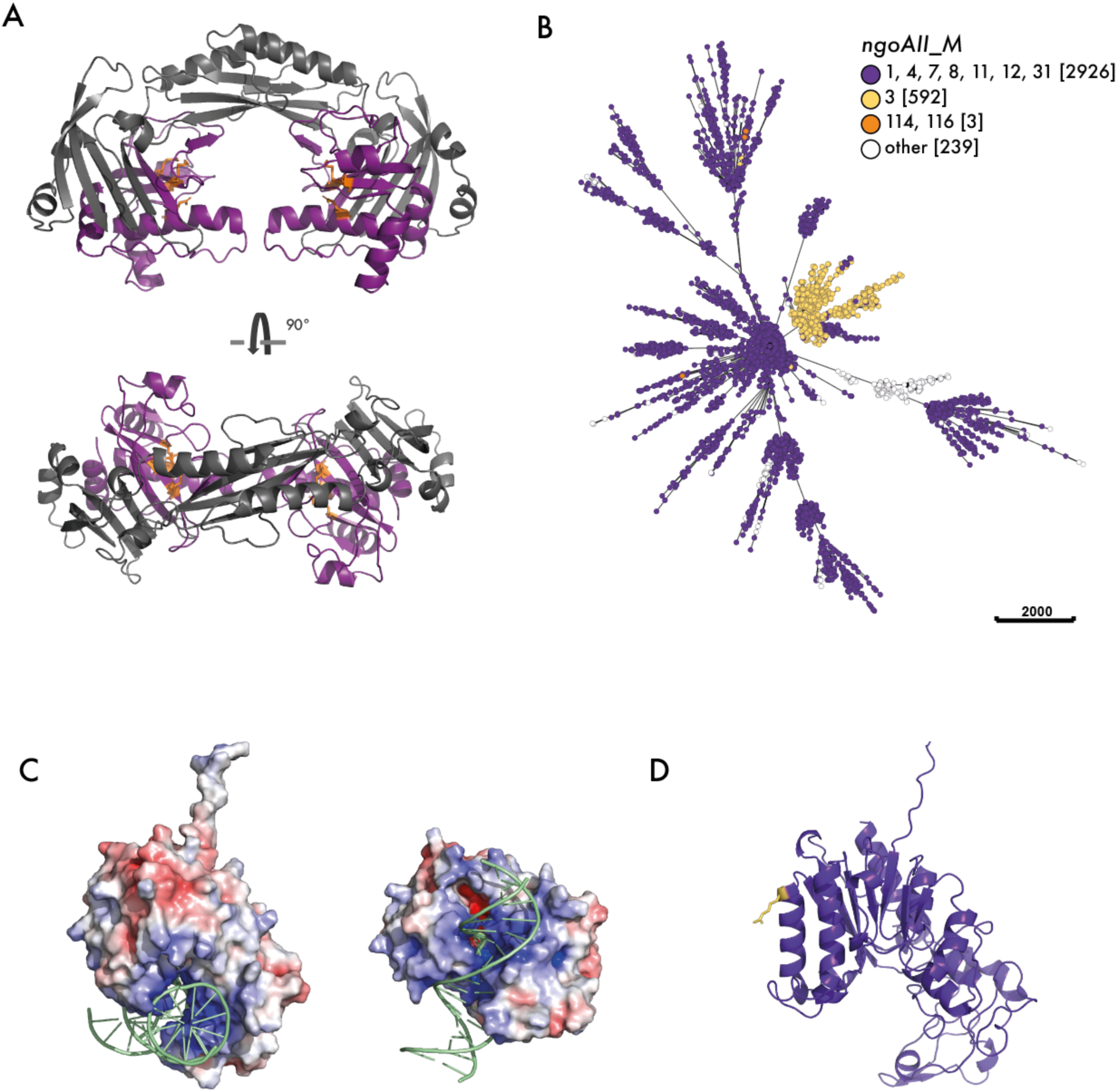
(A) AlphaFold3 prediction of dimeric NgoAII_E (ipTM = 0.76) with structure elements absent from NgoAII_E134 depicted in grey. (B) Minimum spanning tree of 3,761 isolates that were clustered according to core genomic allelic differences (cgMLST v1). Individual isolates are represented as dots that are coloured according to the NgoAII_E variant (A); ngoAII_M alleles encoding identical protein sequences are shown in the same colour. (C) APBS surface charge predictions of NgoAII_M. Regions with positive surface charge are indicated in blue, electronegative regions in red. (D) NgoAII_M with the G145R substitution is indicated in yellow.

**Supplementary Figure 3:**
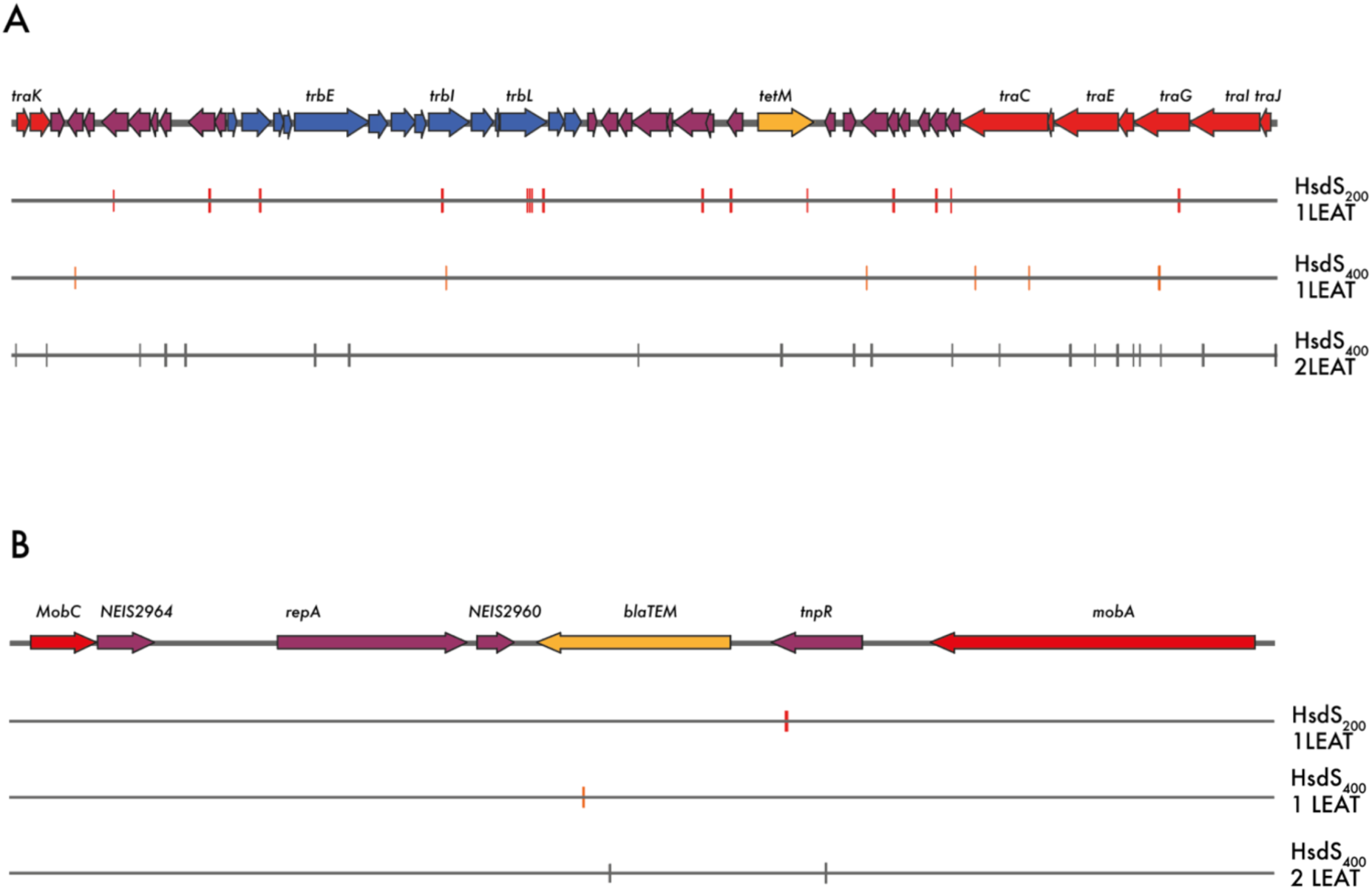
Plasmid maps of (A) pConj and (B) pbla, linearised at their oriTs. Methylation sites of the different HsdS variants are indicated below.

**Supplementary Figure 4:**
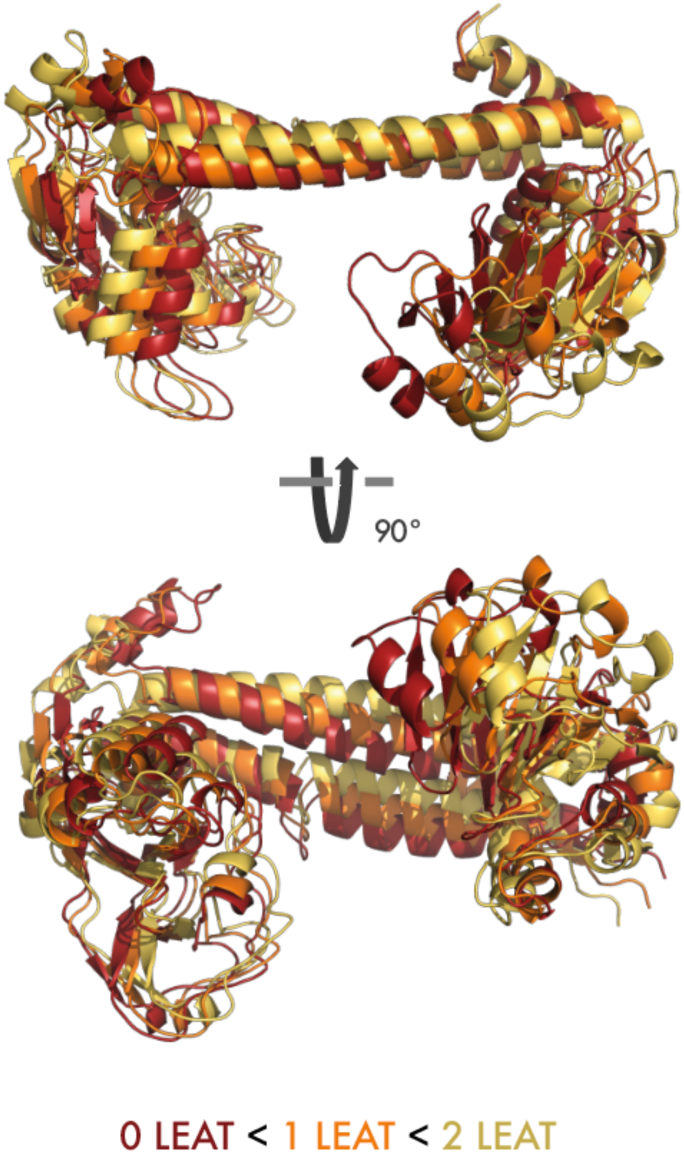
Superimposed Alphafold 3 models of HsdS400 with 0 (red), 1 (orange) and 2 (yellow) LEATs.

**Supplementary Figure 5:**
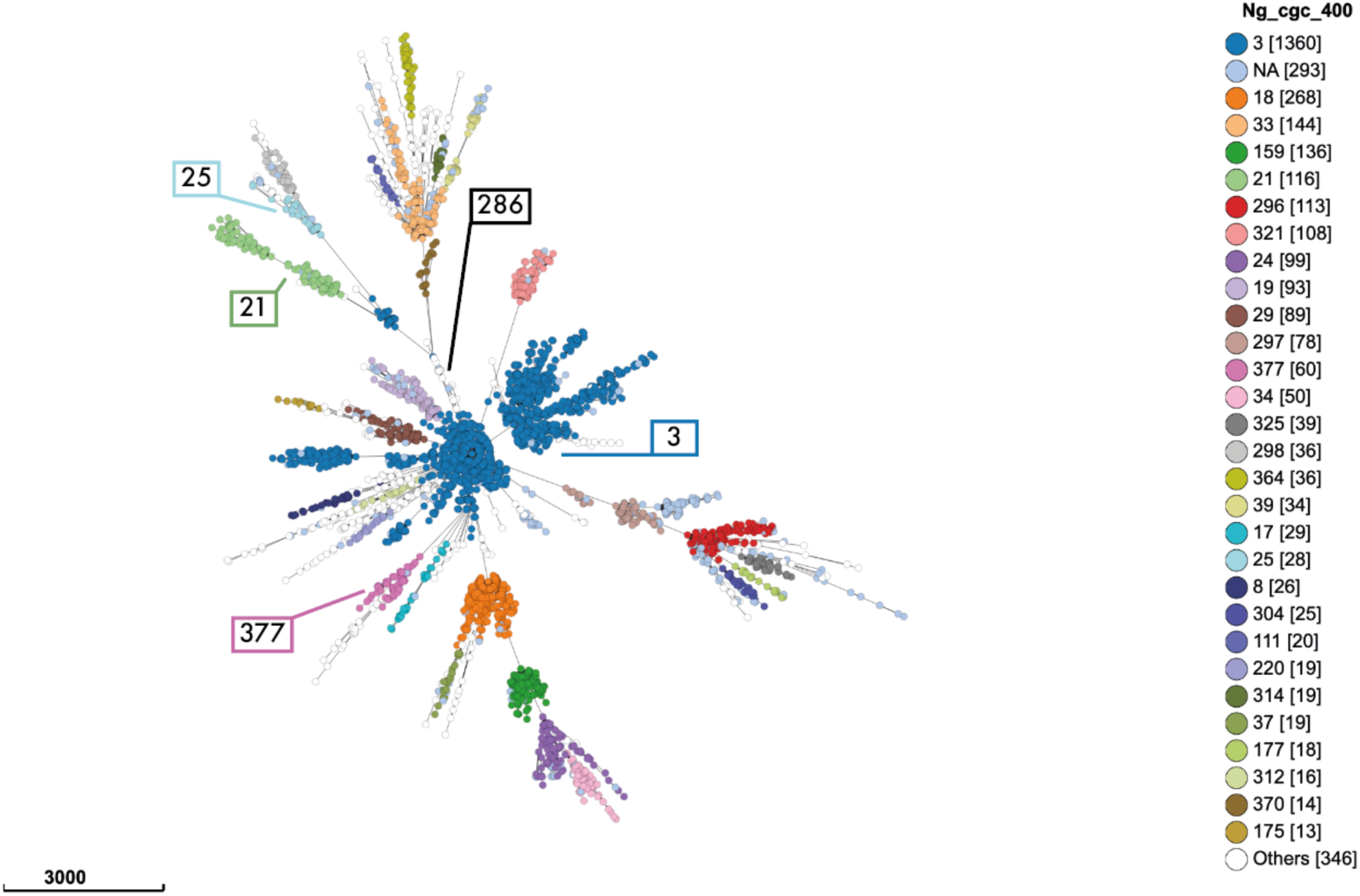
Minimum spanning tree of 3,761 gonococcal isolates clustered by core genome allelic differences (cgMLST v1), with isolates coloured according to their Ng_cgc400. Ng_cgc400s represented by isolates in the mixed strain matings are indicated.

